# *Mycobacterium tuberculosis* – Macrophage Interactome: Molecular Network Structure and Resilience against Antibiotics

**DOI:** 10.1101/2023.04.24.537942

**Authors:** Puente-Mancera Paulina, Valcárcel Antonio, Castillo-Rodal Antonia Isabel, José Díaz

**Author notes:** **Correspondence:** Dra. Antonia Isabel Castillo-Rodal, Dr. José Díaz.

## Abstract

Resistance to several antibiotics against *Mycobacterium tuberculosis* is a serious problem to be solved worldwide. In the present work, we made the statistical analysis of the gene regulatory network of *Mycobacterium tuberculosis* and of the *Mycobacterium tuberculosis*-macrophage interactome to find the probable cause of this resistance. The results from this analysis show that both the gene regulatory network and the interactoma have a hierarchical free scale modular structure that assures a high degree of resilience of these networks against external perturbations. In particular, the interactome is a complex hybrid network that results from the formation of novel links between the *Mycobacterium tuberculosis* and macrophage proteins and from the modification of the previously existing links between the native macrophage proteins, which give rise to novel negative and positive feedback loops that modify the dynamical behavior of the interactome and protect the mycobacterium against the attack with antibiotics by taking control of the macrophage immune response and apoptosis. The statistical analysis of the interactome shows that the highly connected mycobacterium proteins inhA, ahpC, kasA, katG and rpsL exert this control by creating new links with the host proteins FAS and NF-κB. These new hybrid circuits embedded in the hierarchical scale-free modular molecular structure of the interactome produce its high resistance to external perturbations like antibiotics. As consequence, the present work proposes the hypothesis that *Mycobacterium tuberculosis* antibiotic resistance *in vivo* during chronic tuberculosis is only a particular case of a more complex problem that is the interactome *resilience* against antibiotics. Thus, new strategies of drug design are necessary to shatter the complex structure of the *Mycobacterium tuberculosis*-macrophage interactome.

## 1 Introduction

Tuberculosis is an infectious – contagious disease caused by the bacillus *Mycobacterium tuberculosis* (Mtb), the second cause of death by a single infectious agent. About 10.6 million people in the world acquired this disease in 2021, from which 1.6 million people died, and 450,000 became resistant to rifampicin (RR) and isoniazid (INH) call multidrug resistance (MDR). In addition, there are reports of resistance to fluoroquinolones and amikacin, the tuberculosis second-line of treatment (WHO, 2022).

Mtb resistance to antibiotics occurs naturally and intrinsically in different ways (Singh R 2019). However, all the sites of mechanisms of action of antibiotics that produce Mtb resistance are still unknown (Gómez-Tangarife et al., 2018). The emergence of resistance to various antibiotics against tuberculosis raises the need to search for new targets for current and new antibiotics. Thus, it is necessary to know the interactions between Mtb and macrophage host proteins (interactome), and between current and new antibiotics with Mtb proteins and genes.

An alternative tool to analyze the probable resistance to antibiotics or new targets for their identification is network theory where scattered experimental data from interconnected cellular processes are integrated (Sable and Jois, 2015; Shin et al, 2017; Schwab et al, 2020; Breitling, 2010). The theoretical approach to biological networks structure and function allows the integration of disperse experimental data in a coherent model of the spatio-temporal dynamics of interconnected cellular processes.

In this sense, the application of network theory to the analysis of Mbt-host cell interaction in tuberculosis is a guide for the design and use of antibiotics. However, there is a lack of the quantitative information necessary to propose Ordinary Differential Equation (ODEs)-based continuous models for the analysis of the spatio-temporal dynamics of the Mbt infection. In contrast, in the literature (Verma et al, 2021) and in data bases like UNIPROT (https://www.uniprot.org/uniprot/P9WH63; see Model section) there is abundant information about the Mtb-macrophage interactome to allow the construction of the its equivalent network and its statistical analysis. Results from this analysis give up clues about the molecular structure of the Mtb interactome and the mechanisms of antibiotic resistance.

In this form, the objective of this brief communication is to publish the results obtained from the statistical analysis of the Mtb gene regulatory network (GRN) and Mtb-macrophage interactome, which suggest that both the Mtb GRN and the interactome have a hierarchical scale-free modular molecular structure with a set of highly connected bacterial genes and proteins (hubs) that give rise to new molecular hybrid circuits that control the immune response of the macrophage and the entrance to apoptosis propitiating the tuberculosis chronic state. These new hybrid circuits embedded in the hierarchical scale-free modular molecular structure of the interactome produce its high resistance to external perturbations like antibiotics. As consequence, the present work proposes the hypothesis that Mtb antibiotic resistance *in vivo* is only a particular case of a more complex problem that is the interactome *resilience* to antibiotics. Thus, *new strategies of drug design are necessary to shatter the complex structure of the Mtb-macrophage interactome instead of the actual strategies based on antibiotics directed only against punctual Mtb nodes*.

## 2 Methods

Mtb GRN and Mtb-macrophage interactome undirected network were built from the data reported in literature (Singh et al, 2019; Verma et al., 2021; Singh et al., 2021) and in data bases like UNIPROT, TBDreaMDB, GMTVD and MUBII-TB-DB. Gephi 0.9.4 was used to construct the networks and draw them. Mtb GRN graphical representation was made with the Fruchterman-Reingold algorithm (Fruchterman and Reingold, 1991) (Figure 1a). Statistical analysis of the networks was made with Gephi and Matlab R2014a.

**Figure 1.**
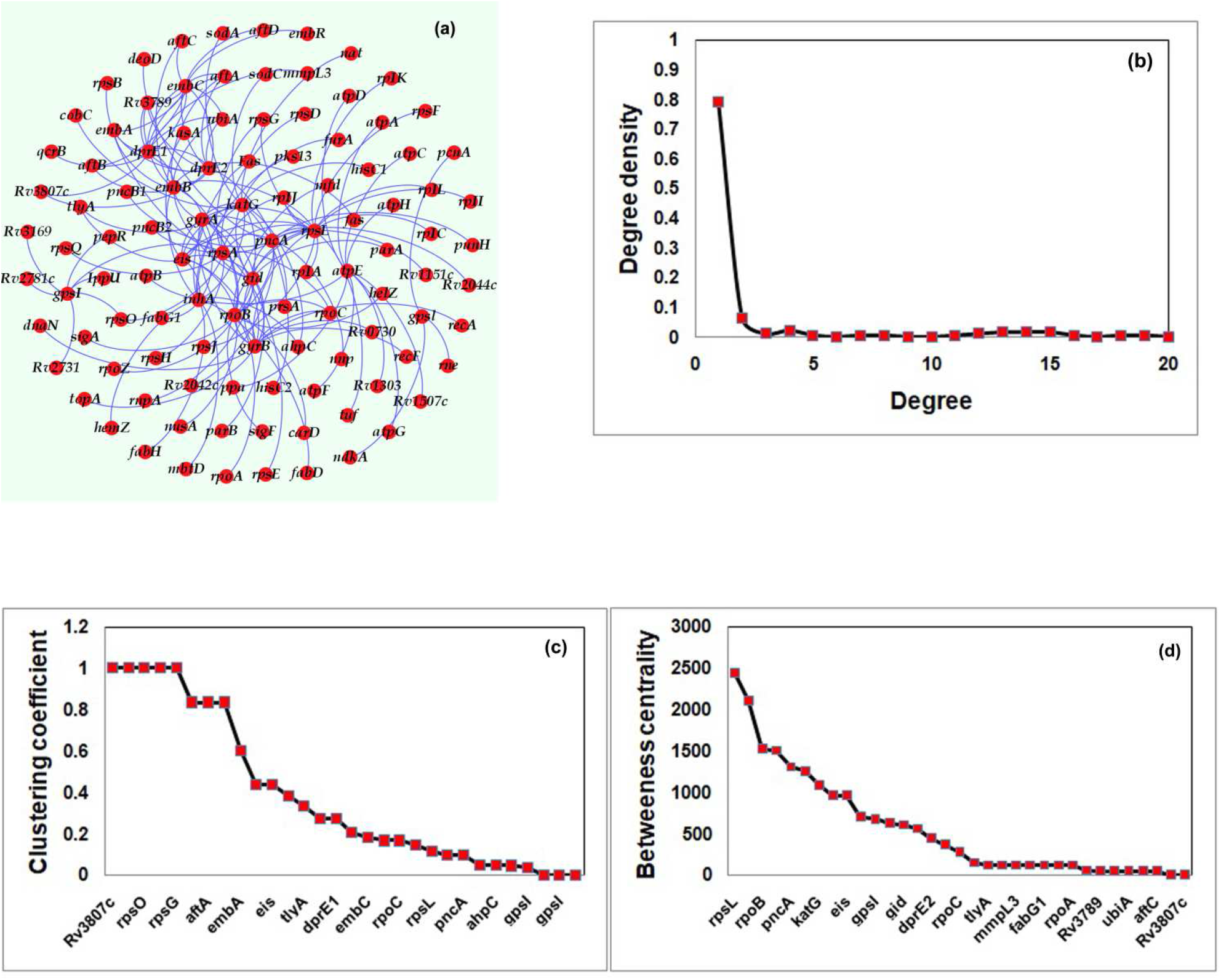
Basic statistical properties of Mtb GRN. (a) Network representation of Mtb GRN was done with the Fruchterman-Reingold algorithm. Red circles represent Mtb genes. Blue lines represent the undirected links. The most connected gene is *rpsL*. (b) Degree distribution of the genes of the Mtb GRN. Y-axis indicates the number of links per gene (degree density), and the x-axis are the genes (nodes) of the network. The degree distribution fits a decaying exponential. The distribution follows the power-law *P* (*k*) ∼ *k* ^−3.679^ with 0.95 of confidence. (c) Clustering coefficient distribution of the network. The y-axis represents the clustering coefficient value indicated and the x-axis represents the corresponding node. A high number or nodes has a clustering coefficient of 0, which indicates a GRN structure with low density of connections. d) Betweenness centrality distribution of the GRN, in this graph the genes *rpsL, rpoB, pncA, gatG, eis, gpsL* and *gidB* have the highest values of this parameter indicating that this nodes lie on the path between most of the other nodes of the network and that a great part of the genetic information of the GRN must flow through these nodes.

The basic statistical analysis of the network includes the determination for each node of the value of the following parameters: degree, degree distribution, clustering coefficient, betweenness centrality and modularity class. Supplementary Material Table 1 (SMT 1) shows the results of this statistical analysis. The mathematical definition of these parameters is discussed in Supplementary Material 2 (SM2).

## 3 Results

### 3.1) Mtb GRN

Data obtained from the search in the literature and databases were used to investigate the structure of Mtb GRN involved in antibiotic resistance, which consists of 187 genes connected in a hierarchical organization. We found that in this GRN there is a set of highly connected genes that control the activation state of the poor connected downstream genes (Figures 1a, 1b). The most connected gene is *rpsL* with 19 links, followed by *rpoB* with 18 links, *embB* with 16 links, followed by a set of genes with 11 to 15 links that includes *katG, gyrA*, and *inhA* among others. The results from the calculus of the degree density confirm a hierarchical organization of this GRN with a set of 149 genes with only one link and 15 genes with high degree (Figure 1b). This degree distribution fits to the power-law distribution *P* (*k*) = 0.7899*k* ^−3.679^ with *R*^*2*^ = 0.9978, where *k* is the degree and *P*(*k*) is the degree density (SM 2). In this case, −4.062 < *γ* < −3.297 with 0.95 of confidence, which clearly indicates a free-scale structure of the Mbt GRN, which suggests that genes *rpsL, rpoB, embB, katG, gyrA, inhA, pncA, gyrB, gidB, dprE2, dprE1, embC, rpsA, eis* and *atpE* form a set of hubs that command the *gene expression* of the bacteria and is reasonable to assume that their proteins have a central role in antibiotic resistance.

A particular characteristic of this GRN is that most of its genes have a small clustering coefficient (Figure 1c) indicating that their neighbors are scarcely connected between them. Only genes like *Rv3789, aftA, ubiA, embA, rpoC, ahpC, tyla, rspG, rplJ, rspO*, and *Rv3807c* have a clustering coefficient 1, which indicates that all the genes in their neighborhood are linked. Furthermore, ∼ 85% of the genes have a clustering coefficient 0 indicating that their neighbors have not links between them. Genes like *gidB, rpoA, kasA, rplC* and *sigA*, among others, belong to this group (Figure 1c).

Finally, the average clustering coefficient of the network (Eq. (4) in SM 2) is 0.313, which corresponds to a network with a small density of connections between the neighbors of each node.

The Mbt GRN has a modularity value of 0.728 (Eq (5) of SM 2), which is far from the negative value for a random network. The statistical analysis of the GRN shows that the network has 34 clusters with sizes that range from 0 to 34 nodes, indicating that the number of nodes distributed between the 34 clusters is larger than the expected number for a random network.

The values of betweeness centrality (Eq. (6) of SM 2) suggest that the Mtb GRN is not a random network but has a hierarchical structure. In fact, genes *rpsL, embB, rspA, rpoB, inhA, katG, pncA, gyrA* and *eis* has the highest values of this parameter indicating that this nodes lie on the path between most of the other nodes of the network (see SM 2) and that great part of the genetic information of the GRN must flow through these nodes (Figure 1d).

### 3.2) Mtb-Macrophage interactome

The Mtb-macrophage interactome consists of 280 linked proteins (Figure 2a) in which the most connected gene are the macrophage proteins RPS3A, RPS8, EEF182, and RPL15 with 26 links, followed by a set of Mtb and macrophage proteins with 15 to 25 links that includes the bacterial proteins pknG, rpsL, kasA and ahpC among others (SMT 1). The results from the calculus of the degree density confirm a hierarchical organization of the interactome with a set of 168 proteins with only one link and 9 proteins with degree 20 o more (Figure 2b). This degree distribution fits to the power-law distribution *P* (*k*) = 0.6032*k* ^−2.945^ with *R*^*2*^ = 0.9871. In this case, −3.391 < *γ* < −2.498 with 0.95 of confidence, which clearly indicates a hierarchical free-scale structure of the interactome. Thus, is reasonable to assume that this particular organization is related to a high resilience of the interactome to the attack with antibiotics (Díaz, 2020; Liu et al., 2022).

**Figure 2.**
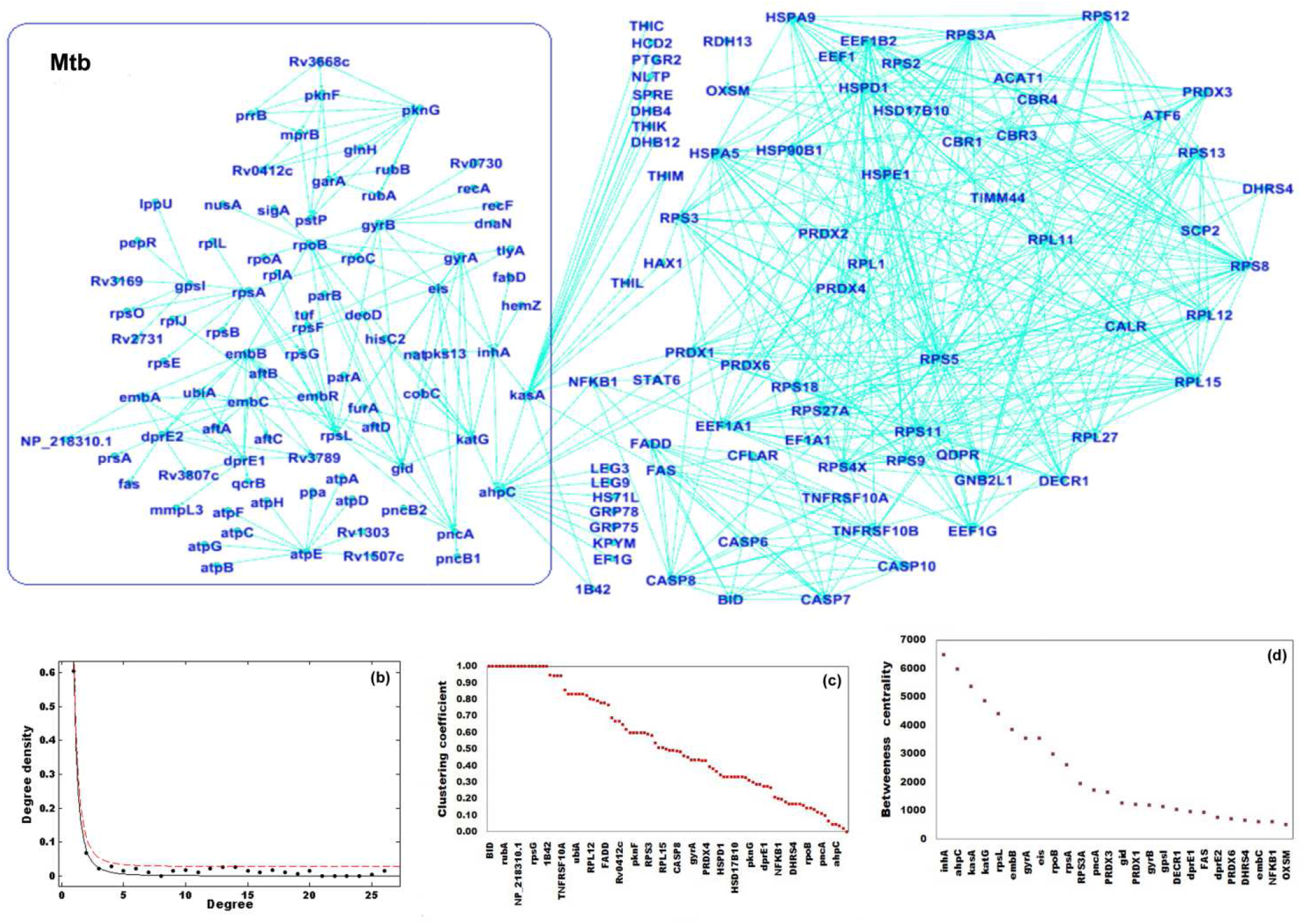
Basic statistical properties of Mbt-Macrophage interactome. (a) Network representation of Mbt-macrophage interactome. Blue lines represent the undirected links. (b) Degree distribution of the 280 proteins of the interactome. Y-axis indicates the number of links per gene (degree density), and the x-axis is the number of nodes with the corresponding degree. The degree distribution fits a decaying exponential. The distribution follows the power-law *P*(*k*) ∼ *k*^−2.945^ with 0.95 of confidence (red line). (c) Clustering coefficient distribution of the network. The y-axis represents the clustering coefficient value indicated and the x-axis represents the corresponding node. A high number or nodes has a clustering coefficient < 1, which indicates an interactome structure with low density of connections. d) Betweenness centrality distribution of the interactome, in this graph host proteins like FAS and NF-κB1 and Mtb proteins like inhA and ahpC have the highest values of this parameter indicating that these nodes lie on the path between most of the other nodes of the network and concentrate great part of the flow of information in the interactome.

Another particular characteristic of the interactome is that most of its proteins have a small clustering coefficient (Figure 2c) indicating that their neighbors are scarcely connected between them.

However, host proteins like BID, CFLAR, RPS2, CBR3, STAT6, and Mtb proteins like Rv3668c, rubA, rub, rsp0, rplJ, among others (see SMT 1), have a clustering coefficient of 1, which indicates that all the proteins in their neighborhood are linked, forming tightly knit groups of proteins.

Furthermore, ∼ 64% of the interactome proteins have a clustering coefficient 0 indicating that their neighbors have not links between them. Proteins like LEG3, STAB1, fabD and atpA belong to this group (Figure 2c and SMT 1). Finally, the average clustering coefficient of the network is 0.19, which corresponds to a network with a small density of connections between the neighbors of each node.

The interactome has a modularity value of 0.628, which is far from the negative value of a random network. The network has 37 clusters with sizes that range from 0 to 37 nodes (SMT 1), indicating that the number of nodes distributed between the 37 clusters is larger than the expected number for a random network, i.e., the interactome has a modular structure.

The values of betweeness centrality presented in SMT I suggest that the interactome is not a random network but has a hierarchical structure. In fact, the Mtb proteins inhA, ahpC, kasA, katG and rpsL and host proteins like FAS and NF-κB1 have the highest values of this parameter indicating that these nodes lie on the path between most of the other nodes of the network and that they concentrate great part of the information that flow through the interactome during the Mtb infection (Figure 2d).

Mtb interacts with the macrophage at the protein level with only two bacterial proteins: ahpC and KasA (Verma et al, 2021). In the interactome, protein ahpC has a degree of 17, a Betweenness centrality value of 5980 and a clustering coefficient of 0.04 (SMT 1). KasA has a degree of 18, a betweenness centrality value of 5369 and a clustering coefficient of 0.02 (SMT 1). Betweenness centrality and clustering coefficient values of these nodes indicate that although the nodes around them are poor connected a great part of the flow of information between the Mtb and the macrophage GRNs is concentrated by ahpC and KasA. Furthermore, ahpC and KasA are linked to proteins pncA and ndh in a closed circuit in such form that any change in the activity of one of these proteins alters the functioning of the circuit (Figure 3). SMT 1 and shows that pncA has a betweenness centrality value of 1710 while ndh has a value of 0 indicating that pncA has an important role in the flow of information from the Mtb GRN to the macrophage GRN (Díaz and Martínez-Mekler, 2022) and probably controls ahpC and KasA activity in the interactome, while ndh has not an important role in this flow of information and is only downstream target of the other three proteins.

**Figure 3.**
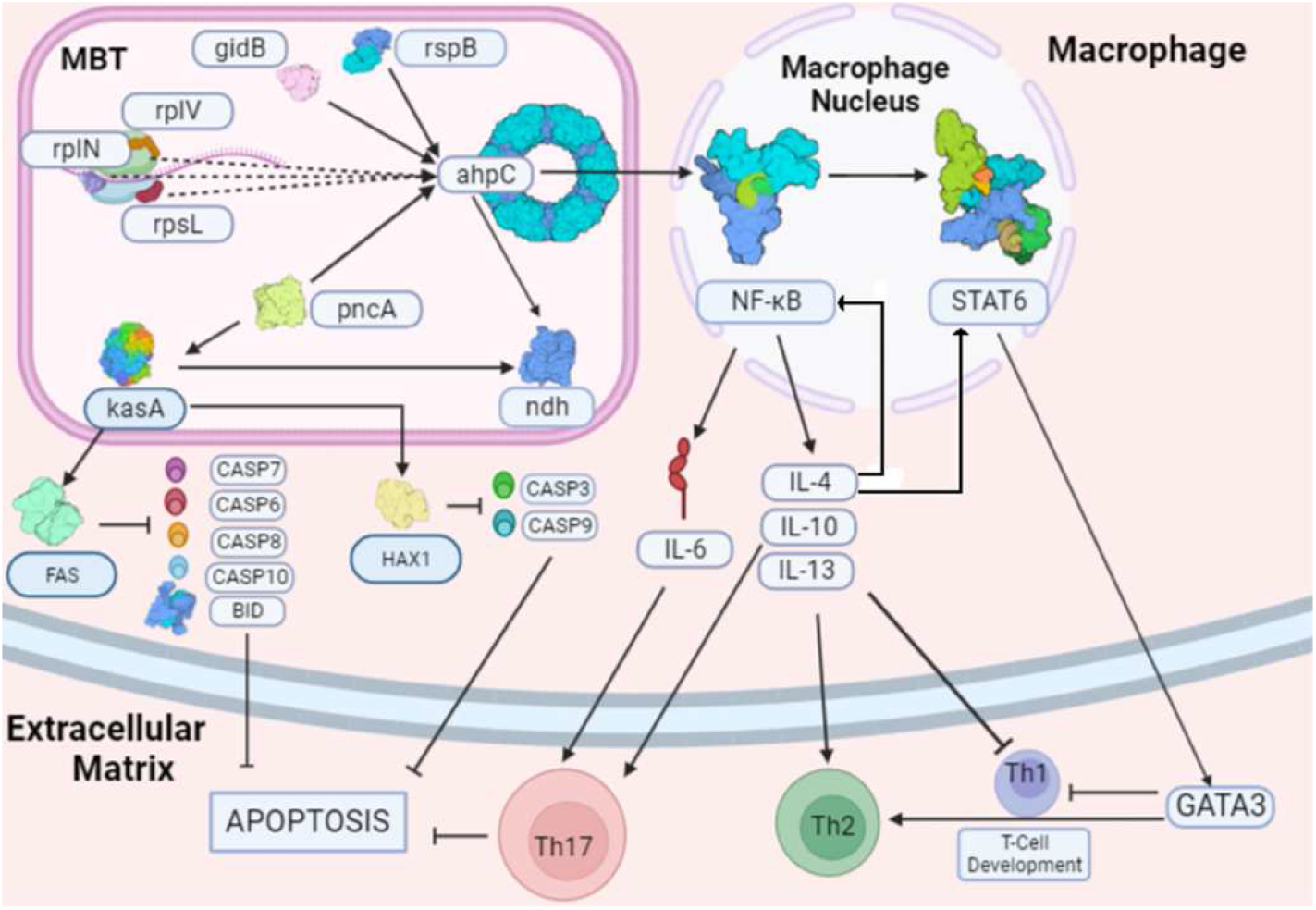
Control of macrophage immune response and apoptosis by *Mycobacterium tuberculosis*. The Mtb-macrophage interactome modifies the host immune response and apoptosis by forming new feedback circuits that dampen the destructive effects of external perturbations like antibiotics. a) Mtb proteins rspL, rspB, rsplv, gidB, and pncA modulate the state of activation of protein ahpC, which turn on the macrophage immune response by the activation of NF kB1, STAT6, GATA3, TGFβ, and IL-6 production. The maturation of Th17 cells induced by IL-6 and TGFβ together with the maturation of Th2 cells induced by GATA3 produces an excess of cytokines Th17, which inhibits apoptosis. IL-4 has two positive feedback circuits with NF kB1 and STAT6; b) Mtb protein pncA modulates the state of activation of KasA which in turn activates the host protein FAS that inhibit apoptosis by caspases 6, 7, 8, and 10; KasA also activates host protein HAX1, which inhibit apoptosis by inhibiting caspases 3 and 9. Protein pncA links ahpC and KasA.

## 4 Discussion

It is clear from the results that the fusion of the Mtb and macrophage GRNs by and interactome gives rise to a complex hybrid network with emergent properties due to the formation of novel links and the modification of the previously existing ones. As consequence, new feedback loops that modify the dynamical behavior of the network are created, assuring a high degree of resilience of the interactome against external perturbations like antibiotics, which is a characteristic of networks with hierarchical modular free-scale structure (Liu et al., 2022). In this form, resistance to antibiotics is only part of a more general problem that is the resilience against antibiotics, in which the molecular structure of the Mtb-macrophage interactome must be broken to defeat the chronic state of tuberculosis.

An important part of the natural resilience of the Mtb-macrophage interactome is due mainly to two related mechanisms: a) the control of the immune response of the macrophage and auxiliary cells and b) the control of macrophage apoptosis. The control of both mechanisms avoids the destruction of the mycobacterium by the natural defense processes of the host cell. It is of interest that both mechanisms are linked to Mtb proteins pncA, KasA and ahpC (Figure 3) (Verma et al., 2021).

In the first case, pncA modulates the state of activation of ahpC giving rise to a host immune response based on the activation of NF-kB1 modulating the activation of the downstream effector protein STAT6 (Figure 3) (Shen and Stavnezer, 1998; Verma et al, 2021). STAT6 promotes IL-4 secretion (Shen and Stavnezer, 1998; Karpathiou et al., 2021), a Th2 phenotype that in excess inhibits the Th1 micromicidal action (Lazarski et al., 2013), and stimulates the activation of GATA3, a regulator of Th2-cell development (Figure 3) (Maier et al., 2013). NF kB1 also activates both macrophage IL-6 production (Bensussen et al., 2021) and the TGFβ signalling pathway (Saas P 2012) (Figure 3). TGFβ is changed from its latent to its active form by dendritic cells (Khalil, 1999), and both IL-6 and activated TGFβ induce the differentiation of Th17A cells and the subsequent stimulation of Th17A phenotype (Figure 3) (Tesmer et al, 2008). In this form, Mbt gene ahpC and its protein ahpC control the immune response of infected macrophage cells. As shown in Figure 3, this control process is also dynamically modulated by proteins rpsL, rspB, rplv, and gidB (SMT 1), which points out that the state of activation of the immune macrophage response is continuously adjusted to damp the effect of internal and external perturbations, like antibiotics, that can disrupt the chronic state of the infection.

In the second case, maturation of Th17A cells induced by IL-6 and TGFβ together with the maturation of Th2 cells induced by GATA3 produces an excess of Th17 cytokines. IL-4 inhibits apoptosis by itself (Figure 3) and forms positive feedback loops with NF kB1 and STAT6, which can lead also to the excess of Th17 interleukins and the exhaustion of Th17A cells causing apoptosis (Boonefoy S 2011, Shen and Chen, 2018). Protein pncA also activates KasA that targets FAS inhibiting the pro-apoptotic caspases 6, 7, 8, and 10 and protein BID (Figure 3) (Verma et al., 2021); In Mbt GRN and Mbt-macrophage interactome KasA also activates the host protein HAX1, which inhibits apoptosis by inhibiting caspases 3 and 9 (Figure 3). In consequence, apoptosis is also dynamically regulated protecting the Mtb from elimination by host cell death. Both mechanisms can explain the resilience of the Mbt GRN and Mbt-macrophage interactome against classical antibiotics, which attack punctually some targets but cannot annihilate the hierarchical modular free-scale molecular structure of both networks that can adjust their functioning and continue working even without some hubs connected like KasA and ahpC, by an unknown mechanism

As consequence, the present work proposes the hypothesis that MBT antibiotic resistance *in vivo* is only a particular case of a more complex problem that is the interactome *resilience* to antibiotics.

Thus, *new strategies of drug design to shatter the complex structure of the Mtb-macrophage interactome are necessary instead of the actual strategies based on antibiotics directed only against particular Mtb nodes*.

## 5 Conclusions

The statistical analysis of Mtb GRN and Mtb-macrophage interactome show that both nets have a hierarchical free-scale modular structure with 15 highly connected bacterial genes that regulate the genetic processes of the Mbt GRN and macrophage function. The most connected gene is *rpsL* followed by *rpoB* and *embB* all associated with streptomycin resistance as well as *gidB*. a Betweenness centrality analysis of the interactome show a small group of Mbt genes and some host proteins with high values of this parameter that concentrate the flow of information in the interactome inducing an anti-inflammatory phenotype in the macrophage and the inhibition of apoptosis due to the formation of novel positive feedback circuits that sustain these phenotypes during chronic tuberculosis producing the high resilience of the disease against the antibiotics. The search for new antibiotics for the treatment of chronic tuberculosis should not be directed only to highly connected genes, whose mutation rate is very high, but also to the most conserved genes with low rate of mutation to avoid higher rates of antibiotic resistance and to be able of break the molecular structure of the interactome.

## Supporting information

SMT1

## 7 Author Contributions

*All authors contributed equally to this work*

## 9 Acknowledgments

JD thanks Erika Juárez Luna for logistical support and also thanks Dra. Elizabeth Castillo Villanueva, de la Facultad de Medicina de la UNAM, for interesting discussions.

