## Supplementary material for "*Mycobacterium tuberculosis* – Macrophage Interactome: Molecular Network Structure and Resilience against Antibiotics": SM2

#### *Supplementary Material 2*

### Network Parameters of the MBT-Macrophage Interactome

Basic statistical analysis of a network includes the calculus of the value of the following parameters for each node: degree, degree distribution, clustering coefficient, average clustering coefficient, betweeness centrality, and modularity class.

The number of links of a node $N_i$ to its neighbor nodes defines its degree $k_i$. The degree distribution of an undirected network is defined as the number of nodes with degree $k$ ($m_k$) divided by the total number of nodes $m$:

$$P(k) = \frac{m_k}{m} \qquad (1)$$

$P(k)$ is a probability distribution where $k$ = 0, 1, 2,…and $\sum_k P(k) = 1$. In random networks $P(k)$ is a binomial distribution, while in scale-free networks $P(k)$ is an decaying exponential. In the last case, $P(k)$ obeys an power-law distribution:

$$P(k) \sim k^{-\gamma} \qquad (2)$$

where the exponent $\gamma$has a value between 2 and 3. This power-law property is independent of the size (scale) of the network and indicates that a few nodes or hubs determine the connectivity of the network, establishing a hierarchical form of organization.

For a node $N_i$ with $l$ links between its neighbors in an undirected graph the clustering coefficient is defined as:

$$C_i = \frac{2l}{k_i(k_i - 1)} \qquad (3)$$

where $C_i$ represents the density of links associated with the node $N_i$, i.e., the proportion of links between the node $N_i$ and its neighbors divided by the number of links that could possibly exist between the neighbors.

The average clustering coefficient of the network with $m$ nodes is defined as:

$$C_N = \frac{1}{m}\sum_{i=1}^{m} C_i \qquad (4)$$

which is simply the average of the clustering coefficient of each node $N_i$.

Modularity is the fraction of links that fall within a cluster minus the expected fraction if links were distributed at random. Modularity indicates the nodes that are more densely connected between them than with the rest, and reveals clues about **the structure** and the **vulnerable spots of a network.** Modularity of a nonrandom network has a value between 0 and 1. The software Gephi uses the Louvain method for community detection:

$$Q = \frac{1}{2m}\sum_{i,j}\left[A_{ij} - \frac{k_i k_j}{2m}\right]\delta\left(c_i, c_j\right) \qquad (5)$$

where $Q$ is the modularity of the network, $A_{ij}$ is the weight of the link between node $i$ and node $j$, $k_i = \sum_j A_{ij}$ are the weights of the links attached to node $i$, $c_i$ is the cluster to which node $i$ is assigned, and $m = \frac{1}{2}\sum_{i,j} Aij$ . In Eq.(5), $\delta(c_i, c_j) = 1$ si $c_j = c_j$, and 0 otherwise. Gephi find out the modularity classes of a network with an algorithm that compares the number of links *among clusters* to the number of links expected in a random network.

Betweeness centrality measures the extent to which a node lies on paths between other nodes by determining all the shortest paths (geodesic paths) between every pairs of nodes, and then counting how many times a node is on a shortest path between two others. If node $i$ lie on the geodesic path from node $s$ to node $t$ then the variable $n_{st}^i$ is 1 when node $i$ lies in the path between $s$ an $t$ and 0 otherwise. Then the betweeness centrality $x_i$ of node $i$ is given by:

$$x_i = \sum_{st} n_{st}^i \qquad (6)$$
