## Supplementary material for "*Mycobacterium tuberculosis* – Macrophage Interactome: Molecular Network Structure and Resilience against Antibiotics": SMT1

### Table 1

### Statistical Analysis of the MBT-Macrophage Interactome

*Macrophage protein names are written in capital letters

| Node | Degree | Clustering | Modularity class | Betweeness Centrality | Antibiotics |
|---|---|---|---|---|---|
| inhA | 14 | 0.06 | 3 | 6473 | Isoniazid |
| ahpC | 17 | 0.04 | 3 | 5980 | Isoniazid |
| kasA | 18 | 0.02 | 3 | 5369 | Isoniazid |
| katG | 15 | 0.10 | 5 | 4857 | Isoniazid |
| rpsL | 19 | 0.12 | 5 | 4388 | Streptomycin |
| embB | 16 | 0.21 | 17 | 3843 | |
| gyrA | 15 | 0.44 | 5 | 3544 | Fluroquinolones |
| eis | 12 | 0.44 | 5 | 3544 | Isoniazid |
| rpoB | 18 | 0.14 | 5 | 2973 | Rifampicin |
| rpsA | 12 | 0.05 | 8 | 2608 | Pyrazinamide |
| RPS3A | 26 | 0.49 | 1 | 1939 | |
| pncA | 14 | 0.11 | 5 | 1710 | Pyrazinamide |
| PRDX3 | 12 | 0.20 | 0 | 1635 | |
| gidB | 13 | 0.38 | 5 | 1272 | Streptomycin |
| PRDX1 | 12 | 0.36 | 0 | 1201 | Isoniazida |
| gyrB | 14 | 0.17 | 5 | 1193 | Fluroquinolones |
| gpsI | 8 | 0.04 | 8 | 1149 | |
| DECR1 | 16 | 0.16 | 0 | 1026 | |

| | | | | | |
|---|---|---|---|---|---|
| dprE1 | 13 | 0.27 | 17 | 951 | Benzothiazinones |
| FAS | 11 | 0.62 | 2 | 935 | |
| dprE2 | 14 | 0.27 | 17 | 761 | Benzothiazinones |
| PRDX6 | 8 | 0.39 | 0 | 706 | |
| DHRS4 | 4 | 0.17 | 0 | 666 | |
| embC | 13 | 0.18 | 17 | 604 | Ethambutol |
| NFKB1 | 5 | 0.20 | 2 | 595 | |
| OXSM | 6 | 0.13 | 0 | 504 | |
| rpoC | 4 | 0.17 | 5 | 448 | |
| RPS11 | 17 | 0.29 | 1 | 269 | |
| embA | 5 | 0.60 | 17 | 268 | |
| HSPA5 | 20 | 0.33 | 1 | 246 | |
| RPS8 | 26 | 0.46 | 1 | 228 | |
| EEF1B2 | 26 | 0.45 | 1 | 221 | |
| ATF6 | 9 | 0.33 | 0 | 218 | |
| CASP8 | 14 | 0.49 | 2 | 205 | |
| tlyA | 3 | 0.33 | 5 | 194 | Rifampicin |
| embR | 2 | 0.00 | 17 | 194 | |
| fabG1 | 2 | 0.00 | 3 | 194 | |
| mmpL3 | 2 | 0.00 | 17 | 194 | |

| | | | | |
|---|---|---|---|---|
| RDH13 | 2 | 0.00 | 0 | 194 |
| rpoA | 2 | 0.00 | 5 | 194 |
| HSPD1 | 13 | 0.35 | 1 | 193 |
| RPL15 | 26 | 0.51 | 1 | 188 |
| TNFRSF10B | 13 | 0.50 | 2 | 172 |
| EEF1A1 | 19 | 0.60 | 1 | 171 |
| CBR1 | 3 | 0.33 | 0 | 165 |
| SCP2 | 7 | 0.14 | 0 | 152 |
| RPS5 | 25 | 0.49 | 1 | 141 |
| RPL11 | 15 | 0.58 | 1 | 137 |
| RPS18 | 11 | 0.51 | 1 | 117 |
| ACAT1 | 6 | 0.27 | 0 | 116 |
| CALR | 10 | 0.29 | 0 | 102 |
| GNB2L1 | 20 | 0.65 | 1 | 101 |
| PRDX2 | 7 | 0.43 | 0 | 101 |
| aftA | 4 | 0.83 | 17 | 88 |
| Rv3789 | 4 | 0.83 | 17 | 88 |
| ubiA | 4 | 0.83 | 17 | 88 |
| aftB | 2 | 0.00 | 17 | 88 |

| | | | | | |
|---|---|---|---|---|---|
| aftC | 2 | 0.00 | 17 | 88 | |
| PRDX4 | 8 | 0.43 | 0 | 87 | |
| RPS3 | 17 | 0.59 | 1 | 80 | |
| HSP90B1 | 12 | 0.48 | 0 | 75 | |
| HAX1 | 2 | 0.00 | 3 | 64 | |
| CBR4 | 4 | 0.17 | 0 | 61 | |
| EEF1 | 8 | 0.54 | 1 | 58 | |
| atpE | 11 | 0.00 | 6 | 55 | Bedaquiline |
| FADD | 10 | 0.78 | 2 | 54 | |
| EEF1G | 13 | 0.44 | 1 | 36 | |
| HSPE1 | 20 | 0.69 | 1 | 28 | |
| CASP7 | 10 | 0.80 | 2 | 25 | |
| pknG | 20 | 0.31 | 37 | 24 | |
| RPS13 | 18 | 0.76 | 1 | 19 | |
| RPS4X | 17 | 0.78 | 1 | 17 | |
| RPL12 | 17 | 0.80 | 1 | 12 | |
| RPS27A | 16 | 0.83 | 1 | 11 | |
| TIMM44 | 5 | 0.30 | 0 | 8 | |
| HSPA9 | 14 | 0.79 | 1 | 7 | |
| HSD17B10 | 4 | 0.33 | 0 | 5 | |

| | | | | |
|---|---|---|---|---|
| RPL27 | 13 | 0.67 | 1 | 5 |
| QDPR | 3 | 0.00 | 0 | 4 |
| RPS12 | 15 | 0.86 | 1 | 3 |
| gpsl | 3 | 0.00 | 7 | 3 |
| pknF | 12 | 0.60 | 37 | 3 |
| garA | 10 | 0.60 | 37 | 2 |
| mprB | 10 | 0.60 | 37 | 2 |
| ddn | 2 | 0.00 | 4 | 1 |
| CASP10 | 9 | 0.94 | 2 | 1 |
| CASP6 | 9 | 0.94 | 2 | 1 |
| TNFRSF10A | 9 | 0.94 | 2 | 1 |
| RPS9 | 14 | 0.95 | 1 | 1 |
| Rv0412c | 6 | 0.67 | 37 | 1 |
| EF1A1 | 3 | 0.33 | 1 | 0 |
| prrB | 8 | 0.83 | 37 | 0 |
| pstP | 8 | 0.83 | 37 | 0 |
| RPL1 | 3 | 0.33 | 1 | 0 |
| BID | 8 | 1.00 | 2 | 0 |
| CFLAR | 7 | 1.00 | 2 | 0 |

| | | | | |
|---|---|---|---|---|
| rubA | 6 | 1.00 | 37 | 0 |
| rubB | 6 | 1.00 | 37 | 0 |
| Rv3668c | 6 | 1.00 | 37 | 0 |
| RPS2 | 5 | 1.00 | 1 | 0 |
| glnH | 4 | 1.00 | 37 | 0 |
| 1B42 | 2 | 1.00 | 3 | 0 |
| CBR3 | 2 | 1.00 | 0 | 0 |
| NP_218310.1 | 2 | 1.00 | 17 | 0 |
| rplJ | 2 | 1.00 | 8 | 0 |
| rpsG | 2 | 1.00 | 8 | 0 |
| rpsO | 2 | 1.00 | 8 | 0 |
| Rv3807c | 2 | 1.00 | 17 | 0 |
| STAT6 | 2 | 1.00 | 2 | 0 |
| THIL | 2 | 1.00 | 3 | 0 |
| THIM | 2 | 1.00 | 3 | 0 |
| aftD | 1 | 0.00 | 17 | 0 |
| ald | 1 | 0.00 | 16 | 0 |
| alrA | 1 | 0.00 | 28 | 0 |
| atpA | 1 | 0.00 | 6 | 0 |
| atpB | 1 | 0.00 | 6 | 0 |

| | | | | |
|---|---|---|---|---|
| atpC | 1 | 0.00 | 6 | 0 |
| atpD | 1 | 0.00 | 6 | 0 |
| atpF | 1 | 0.00 | 6 | 0 |
| atpG | 1 | 0.00 | 6 | 0 |
| atpH | 1 | 0.00 | 6 | 0 |
| BAG6 | 1 | 0.00 | 0 | 0 |
| BLVRB | 1 | 0.00 | 0 | 0 |
| carD | 1 | 0.00 | 5 | 0 |
| clpC | 1 | 0.00 | 15 | 0 |
| cobC | 1 | 0.00 | 5 | 0 |
| cycA | 1 | 0.00 | 23 | 0 |
| deoD | 1 | 0.00 | 5 | 0 |
| dfrA | 1 | 0.00 | 27 | 0 |
| DHB12 | 1 | 0.00 | 3 | 0 |
| DHB4 | 1 | 0.00 | 3 | 0 |
| dnaN | 1 | 0.00 | 5 | 0 |
| EF1G | 1 | 0.00 | 3 | 0 |
| ethA | 1 | 0.00 | 12 | 0 |
| ethR | 1 | 0.00 | 11 | 0 |

| | | | | |
|---|---|---|---|---|
| fabD | 1 | 0.00 | 3 | 0 |
| fabH | 1 | 0.00 | 3 | 0 |
| fadE24 | 1 | 0.00 | 22 | 0 |
| fas | 1 | 0.00 | 17 | 0 |
| Fas | 1 | 0.00 | 3 | 0 |
| fbiA | 1 | 0.00 | 31 | 0 |
| fbiB | 1 | 0.00 | 32 | 0 |
| fbiC | 1 | 0.00 | 19 | 0 |
| fgd1 | 1 | 0.00 | 26 | 0 |
| folC | 1 | 0.00 | 34 | 0 |
| furA | 1 | 0.00 | 5 | 0 |
| gidA | 1 | 0.00 | 9 | 0 |
| GRP75 | 1 | 0.00 | 3 | 0 |
| GRP78 | 1 | 0.00 | 3 | 0 |
| HCD2 | 1 | 0.00 | 3 | 0 |
| helZ | 1 | 0.00 | 5 | 0 |
| hemZ | 1 | 0.00 | 3 | 0 |
| hisC1 | 1 | 0.00 | 5 | 0 |
| hisC2 | 1 | 0.00 | 5 | 0 |
| HS71L | 1 | 0.00 | 3 | 0 |

| | | | | | |
|---|---|---|---|---|---|
| iniA | 1 | 0.00 | 33 | 0 | |
| lppU | 1 | 0.00 | 8 | 0 | |
| KDSR | 1 | 0.00 | 0 | 0 | |
| KPYM | 1 | 0.00 | 3 | 0 | |
| LEG3 | 1 | 0.00 | 3 | 0 | |
| LEG9 | 1 | 0.00 | 3 | 0 | |
| mbtD | 1 | 0.00 | 3 | 0 | |
| mfd | 1 | 0.00 | 5 | 0 | |
| mshA | 1 | 0.00 | 13 | 0 | |
| nat | 1 | 0.00 | 5 | 0 | |
| ndh | 1 | 0.00 | 14 | 0 | |
| ndkA | 1 | 0.00 | 7 | 0 | Isoniazid |
| NLTP | 1 | 0.00 | 3 | 0 | |
| nnp | 1 | 0.00 | 5 | 0 | |
| NP_214519.2 | 1 | 0.00 | 5 | 0 | |
| NP_214520.1 | 1 | 0.00 | 5 | 0 | |
| NP_214720.1 | 1 | 0.00 | 17 | 0 | |
| NP_214856.1 | 1 | 0.00 | 33 | 0 | |
| NP_214921.1 | 1 | 0.00 | 26 | 0 | |

| | | | | |
|---|---|---|---|---|
| NP_215000.1 | 1 | 0.00 | 13 | 0 |
| NP_215002 | 1 | 0.00 | 10 | 0 |
| NP_215181.1 | 1 | 0.00 | 5 | 0 |
| NP_215182.1 | 1 | 0.00 | 5 | 0 |
| NP_215192.1 | 1 | 0.00 | 21 | 0 |
| NP_215196.1 | 1 | 0.00 | 5 | 0 |
| NP_215215.1 | 1 | 0.00 | 24 | 0 |
| NP_215689.1 | 1 | 0.00 | 19 | 0 |
| NP_215783.1 | 1 | 0.00 | 17 | 0 |
| NP_215821.1 | 1 | 0.00 | 6 | 0 |
| NP_215925.1 | 1 | 0.00 | 29 | 0 |
| NP_215999.1 | 1 | 0.00 | 3 | 0 |
| NP_216000.1 | 1 | 0.00 | 3 | 0 |
| NP_216146.1 | 1 | 0.00 | 8 | 0 |
| NP_216210.1 | 1 | 0.00 | 5 | 0 |
| NP_216220.1 | 1 | 0.00 | 23 | 0 |
| NP_216370.1 | 1 | 0.00 | 14 | 0 |
| NP_216424.1 | 1 | 0.00 | 5 | 0 |
| NP_216495.2 | 1 | 0.00 | 36 | 0 |
| NP_216559.1 | 1 | 0.00 | 5 | 0 |

| | | | | |
|---|---|---|---|---|
| NP_216761.1 | 1 | 0.00 | 3 | 0 |
| NP_216932.2 | 1 | 0.00 | 5 | 0 |
| NP_216944.1 | 1 | 0.00 | 3 | 0 |
| NP_216963.1 | 1 | 0.00 | 34 | 0 |
| NP_217051.1 | 1 | 0.00 | 18 | 0 |
| NP_217279.1 | 1 | 0.00 | 27 | 0 |
| NP_217280.1 | 1 | 0.00 | 35 | 0 |
| NP_217296.1 | 1 | 0.00 | 16 | 0 |
| NP_217299.1 | 1 | 0.00 | 8 | 0 |
| NP_217655.1 | 1 | 0.00 | 22 | 0 |
| NP_217778.1 | 1 | 0.00 | 31 | 0 |
| NP_217779.1 | 1 | 0.00 | 32 | 0 |
| NP_217783.1 | 1 | 0.00 | 25 | 0 |
| NP_217940.1 | 1 | 0.00 | 28 | 0 |
| NP_217974.1 | 1 | 0.00 | 5 | 0 |
| NP_218064.1 | 1 | 0.00 | 4 | 0 |
| NP_218118.1 | 1 | 0.00 | 20 | 0 |
| NP_218307.1 | 1 | 0.00 | 17 | 0 |
| NP_218308.1 | 1 | 0.00 | 17 | 0 |

| | | | | |
|---|---|---|---|---|
| NP_218312.1 | 1 | 0.00 | 17 | 0 |
| NP_218371.1 | 1 | 0.00 | 12 | 0 |
| NP_218372.1 | 1 | 0.00 | 11 | 0 |
| NP_218436.2 | 1 | 0.00 | 9 | 0 |
| nusA | 1 | 0.00 | 5 | 0 |
| panD | 1 | 0.00 | 20 | 0 |
| parA | 1 | 0.00 | 5 | 0 |
| parB | 1 | 0.00 | 5 | 0 |
| pcnA | 1 | 0.00 | 7 | 0 |
| pepQ | 1 | 0.00 | 18 | 0 |
| pepR | 1 | 0.00 | 8 | 0 |
| pks13 | 1 | 0.00 | 3 | 0 |
| PLS1 | 1 | 0.00 | 3 | 0 |
| pncB1 | 1 | 0.00 | 5 | 0 |
| pncB2 | 1 | 0.00 | 5 | 0 |
| ppa | 1 | 0.00 | 6 | 0 |
| prsA | 1 | 0.00 | 17 | 0 |
| PTGR2 | 1 | 0.00 | 3 | 0 |
| punH | 1 | 0.00 | 5 | 0 |
| qcrB | 1 | 0.00 | 17 | 0 |

| | | | | |
|---|---|---|---|---|
| recA | 1 | 0.00 | 5 | 0 |
| recF | 1 | 0.00 | 5 | 0 |
| ribD | 1 | 0.00 | 29 | 0 |
| rmlD | 1 | 0.00 | 25 | 0 |
| rne | 1 | 0.00 | 7 | 0 |
| rnpA | 1 | 0.00 | 5 | 0 |
| rpIA | 1 | 0.00 | 5 | 0 |
| rpIC | 1 | 0.00 | 5 | 0 |
| rpII | 1 | 0.00 | 5 | 0 |
| rpIK | 1 | 0.00 | 5 | 0 |
| rpIL | 1 | 0.00 | 5 | 0 |
| rplC | 1 | 0.00 | 24 | 0 |
| rpoZ | 1 | 0.00 | 5 | 0 |
| rpsB | 1 | 0.00 | 8 | 0 |
| rpsD | 1 | 0.00 | 8 | 0 |
| rpsE | 1 | 0.00 | 8 | 0 |
| rpsF | 1 | 0.00 | 5 | 0 |
| rpsH | 1 | 0.00 | 8 | 0 |
| rpsJ | 1 | 0.00 | 5 | 0 |

| | | | | | |
|---|---|---|---|---|---|
| rpsQ | 1 | 0.00 | 8 | 0 | |
| RSP18 | 1 | 0.00 | 0 | 0 | |
| Rv0488 | 1 | 0.00 | 10 | 0 | |
| Rv0678 | 1 | 0.00 | 21 | 0 | |
| Rv0730 | 1 | 0.00 | 5 | 0 | |
| Rv1151c | 1 | 0.00 | 5 | 0 | |
| Rv1303 | 1 | 0.00 | 6 | 0 | |
| Rv1507c | 1 | 0.00 | 6 | 0 | |
| Rv1979c | 1 | 0.00 | 36 | 0 | |
| Rv2042c | 1 | 0.00 | 5 | 0 | |
| Rv2044c | 1 | 0.00 | 5 | 0 | |
| Rv2731 | 1 | 0.00 | 8 | 0 | |
| Rv2781c | 1 | 0.00 | 8 | 0 | |
| Rv3169 | 1 | 0.00 | 8 | 0 | |
| sigA | 1 | 0.00 | 5 | 0 | |
| sigF | 1 | 0.00 | 5 | 0 | |
| sodA | 1 | 0.00 | 5 | 0 | Isoniazid |
| sodC | 1 | 0.00 | 5 | 0 | |
| SPRE | 1 | 0.00 | 3 | 0 | |
| STAB1 | 1 | 0.00 | 4 | 0 | |

| | | | | |
|---|---|---|---|---|
| THIC | 1 | 0.00 | 3 | 0 |
| THIK | 1 | 0.00 | 3 | 0 |
| thyA | 1 | 0.00 | 35 | 0 |
| topA | 1 | 0.00 | 5 | 0 |
| tuf | 1 | 0.00 | 5 | 0 |
| whiB7 | 1 | 0.00 | 30 | 0 |
| YP_177940.1 | 1 | 0.00 | 30 | 0 |
| YP_177995.1 | 1 | 0.00 | 15 | 0 |
